# The phosphodiesterase-5 inhibitor vardenafil reverses sleep deprivation-induced amnesia in mice

**DOI:** 10.64898/2026.03.24.713921

**Authors:** Camilla Paraciani, Carlo Castoldi, Diana Madalina Popescu, Elroy L. Meijer, Onno Van Den Hoed, Adithya Sarma, Sophia Wilhelm, Nienke de Vries, Linda Maria Requie, Esmee Y.G. D’Costa, Dimitrios Tantis Tapeinos, Pim R.A. Heckman, Ewelina Knapska, Peter Meerlo, Bianca Ambrogina Silva, Robbert Havekes

**Author notes:** Equal contribution (Bianca Ambrogina Silva, Robbert Havekes).

## Abstract

Sleep deprivation (SD) disrupts memory processes, particularly those dependent on the hippocampus. Six hours of SD after training in a hippocampus-dependent task typically induces amnesia in mice and impairs performance upon memory testing later. However, we previously demonstrated that object-location memories (OLMs) encoded under SD conditions can be recovered several days later, suggesting that these memories were not lost but suboptimally stored. Given that engrams of a specific memory are distributed across multiple functionally connected brain regions, we hypothesized that SD-induced amnesia arises from disrupted network alterations extending beyond the hippocampus. Consistent with this, brain-wide cFos mapping revealed a widespread reduction in cFos in memory associated regions during recall in SD mice and connectivity analysis identified the hippocampus as a central hub in this network. Since cGMP signaling modulates memory processes, we next tested whether the cGMP-specific PDE5 inhibitor vardenafil could restore access to these latent memories. One day after training, vardenafil reversed SD-induced OLM impairment when administered 30 minutes before testing, but this effect was lost when testing occurred several days later. To achieve persistent access to OLMs formed under SD conditions, we combined vardenafil treatment with optogenetic engram stimulation. This combined approach successfully maintained OLM retrievability for several days post-manipulation. Crucially, successful retrieval in these mice was associated with a significant increase in engram cell reactivation within the dorsal dentate gyrus compared to mice that failed to recall. Collectively, these findings provide novel insight into the molecular and network mechanisms underlying SD-induced amnesia and offer a strong rationale for developing targeted PDE5-mediated therapies to reverse SD-related memory deficits.

**Highlights:**

- SD-induced amnesia is associated with reduced cFos expression within memory-associative circuitry
- The phosphodiestarase-5-inhibitor vardenafil can be used to restore memory access
- Combining optogenetics with vardenafil treatment sustains memory retrieval over several days
- Successful retrieval reflects increased reactivation of engram cells in the dentate gyrus

## Introduction

Sleep is a fundamental physiological state essential for both mental and physical health, supporting the homeostasis of numerous bodily functions. Among its multiple functions, sleep has long been proposed to play a critical role in long-term memory consolidation^1–3^. Studies have shown that even brief periods of sleep can enhance declarative memory^4,5^, while sleep loss impairs memory in both humans and rodents^6–10^.

Network analyses have shown that the hippocampus is central to memory formation and operates within a distributed network of cortical and subcortical regions. Successful memory consolidation and later recall depend on the coordinated activity of engram ensembles across different brain areas^11–13^. Brain-wide activity mapping^13,14^ provides a powerful approach for assessing the distributed neural basis of memory and associated dysfunctions^15–19^. However, to date such a network analysis has not yet been employed to determine how the distributed neural circuits underlying memory are perturbed by sleep deprivation (SD).

Brief periods of SD after learning disrupt hippocampus-dependent memory consolidation, particularly the consolidation of hippocampus-dependent memories, through dysregulation of key signaling pathways^2,9,20–27^. Specifically, SD compromises cellular processes important for memory consolidation within the hippocampus, including reduced cyclic adenosine monophosphate (cAMP)/PKA signaling, attenuated mTORC1-dependent protein synthesis, altered dendritic spine density, and impaired long-term potentiation^24,28–35^. Collectively, these alterations in synaptic plasticity processes are thought to be the primary cause of SD-induced amnesia.

Recent findings from our laboratory indicate that memories formed under SD are not lost but rather suboptimally stored, since they can be retrieved through optogenetics engram reactivation or pharmacological treatment targeting the cAMP-PKA signalling pathway^28^. These findings highlight that SD impairs memory accessibility rather than erasing stored information, and that targeting cyclic nucleotide signaling can restore recall. Whether similar effects can be observed for the other major cyclic nucleotides remains to be determined. For example, the cyclic guanosine monophosphate (cGMP)-protein kinase G (PKG) pathway is essential for early-phase long-term potentiation (E-LTP) and early consolidation; modulates synaptic efficacy and supports activity-dependent plasticity^36–40^. Previous studies suggest that pharmacological elevation of cGMP, achieved through inhibition of phosphodiesterase-5 (PDE5), which selectively hydrolyzes cGMP, modulates specific aspects of hippocampus-dependent memory formation in rodents^37,41,42^. However, these effects have predominantly been attributed to facilitation of acquisition or consolidation under baseline conditions. In contrast, the contribution of cGMP signaling to memory retrieval remains unknown. No study to date has directly examined whether acute PDE5 inhibition at the time of retrieval can enhance memory expression, nor whether such manipulation can rescue retrieval deficits induced by SD occurring after learning. Given the established involvement of cGMP in hippocampal synaptic plasticity, acquisition and early memory consolidation, we next investigated whether enhancing cGMP signaling could restore object location memory accessibility under SD conditions. Overall, this study aims to provide mechanistic insights into how SD disrupts hippocampus-dependent memory at both molecular and network levels, and to identify potential therapeutic targets within distributed memory networks that may inform strategies to mitigate the cognitive consequences of insufficient sleep.

## Materials and Methods

An extended description of the different experiments can be found in the supplementary materials.

### Animals

All experimental procedures followed national Central Animal Procedures (CCD) and Institutional Animal Welfare Committee (IvD) guidelines, in line with Directive 2010/63/EU. For whole-brain activity mapping (Fig. 1) and pharmacological studies without engram labeling (Fig. S7), adult male C57Bl/6J mice were obtained from Charles River Laboratories at 6 weeks of age, pair-housed upon arrival, and individually housed one week before the experiment. Animals were maintained on a 12-hour light/12-hour dark cycle with ad libitum food and water. All mice were 2 to 5 months old at testing, which was conducted at the beginning of the light phase. For Figure S7, mice were reused once to comply with guidelines for reduction: each animal underwent training and testing twice, but with different objects in different locations, and animals were treated with drugs only once. For the engram-tagging studies (Fig. 2, 3), adult male c-fos-tTA mice (Jackson Labs strain #:018306) on a C57Bl6/J background were bred in our facility, group-housed until surgery, and individually housed afterwards. Animals were kept on standard chow (Altromin), switched to doxycycline (Dox)-supplemented chow (40 mg/kg) one week before surgery, and remained on Dox until the day before training. Twenty-four hours before training, the diet was switched back to standard chow without Dox to allow engram tagging the day after. Immediately after training, mice were returned to high-dose Dox chow (1000 mg/kg; to accelerate the closure of the tagging window) and remained on this diet for the rest of the experiment (*i.e.*, ^28^). For Figures 1, 2, and 3 experiments, animals were euthanized by transcardial perfusion 90 minutes after behavioral testing; for Figure S7, animals were sacrificed by cervical dislocation.

**Figure 1.**
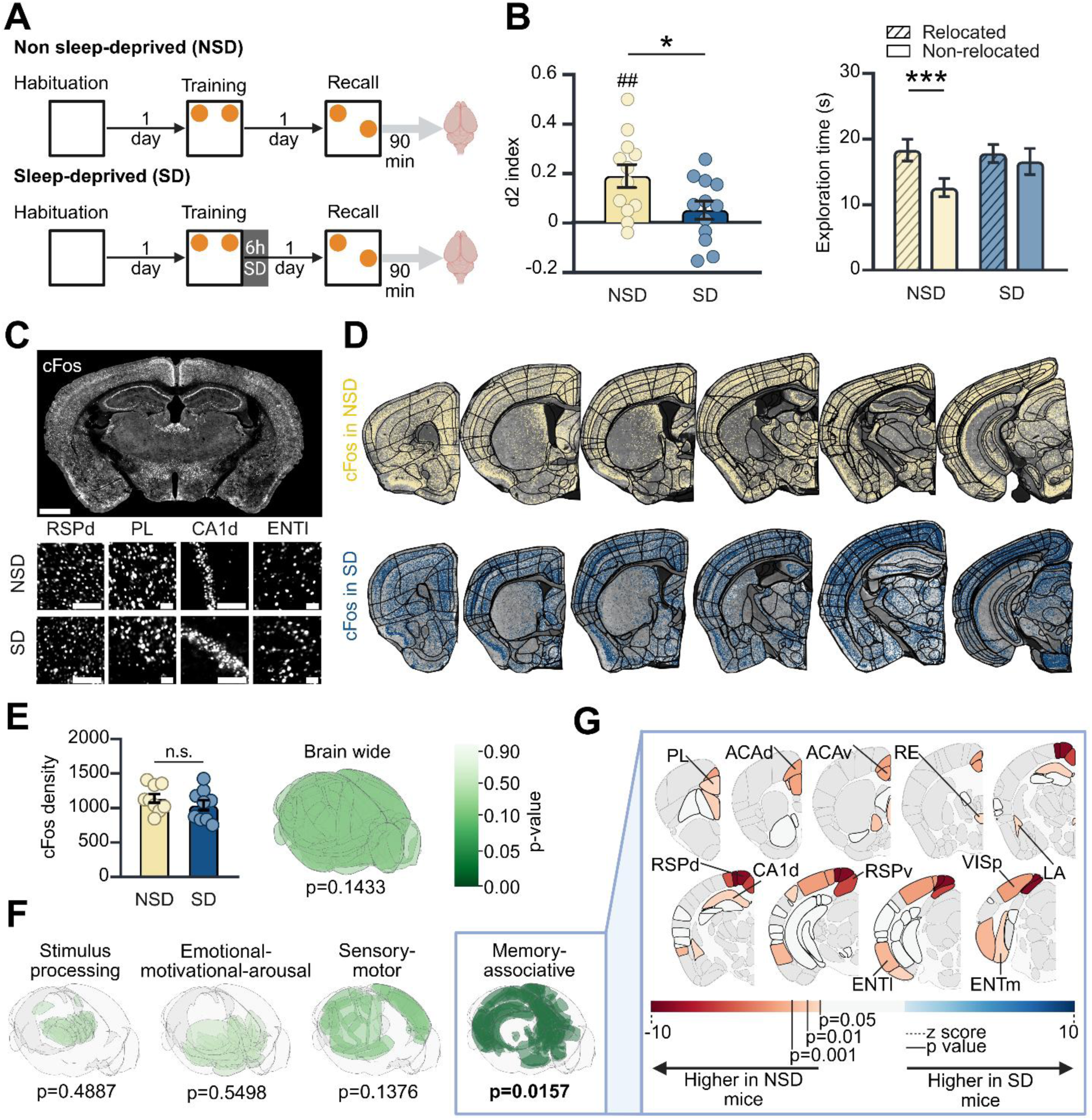
Impaired retrievability to OLMs formed during sleep deprivation is associated with reduced cFos activity in memory circuits. **(A)** Schematic representation of the experimental procedure, n=12 **(B)** Left: d2 index. NSD mice scored above chance, indicating intact memory retrieval, whereas SD mice failed to detect spatial novelty (one-sample t-test: NSD p = 0.0021, SD [N.S.] p = 0.2191), reflecting memory inaccessibility. Right: Exploration times during recall. NSD mice showed a significant preference for the relocated object, whereas SD mice failed to discriminate between objects, reflecting OLM retrievability deficit. Raw values are presented for ease of interpretation; however, statistical analysis for the right panel was performed on log₁₀-transformed data to satisfy normality assumptions. **(C)** Representative images of cFos immunohistochemistry. Top: whole-section representation, scale bar, 1000 μm. Bottom: representative sample regions: RSPd, PL, CA1d and ENTl; scale bar, 100 μm (RSPd, CA1d) and 40μm (PL, ENTl). **(D)** Representative examples of atlas registration and positive cell segmentation. cFos detections are shown for the two conditions: NSD group, yellow; SD group, blue. The black reticulate represents the boundaries of Allen CCFv3’s brain regions, after being mapped onto each brain section with ABBA. **(E)** Brain-wide comparison of cFos density between SD and NSD mice following memory recall. Left: total cFos density (n=9, unpaired t test, P=0.3496). Right: global region by region comparison (n=9 mice, for n=214 regions, mean-centered PLS, p = 0.1433, 10000 permutations). **(F)** cFos density comparison between SD and NSD mice following memory recall in four macrocircuits (mean-centered PLS,10000 permutations): stimulus processing (n=40 regions), sensorimotor function (n=25 regions), emotional–motivational regulation (n=61 regions), and memory–associative (n=46 regions) processes. **(G)** cFos density post-hoc pairwise comparisons between SD and NSD within the memory-associative macrocircuit (n=9 mice for n=46 regions; Supplementary Table 1). Regions in grey are not part of the memory-associative circuit; regions in white are part of the memory-associative circuit but do not significantly differ between the two groups (|z-score| < 1.96). No brain region showed higher cFos expression in SD mice relative to NSD mice. Detailed description of statistics can be found in the Statistical Analysis Table S2. Created using BraiAn, BrainGlobe, and BioRender.com.

**Figure 2.**
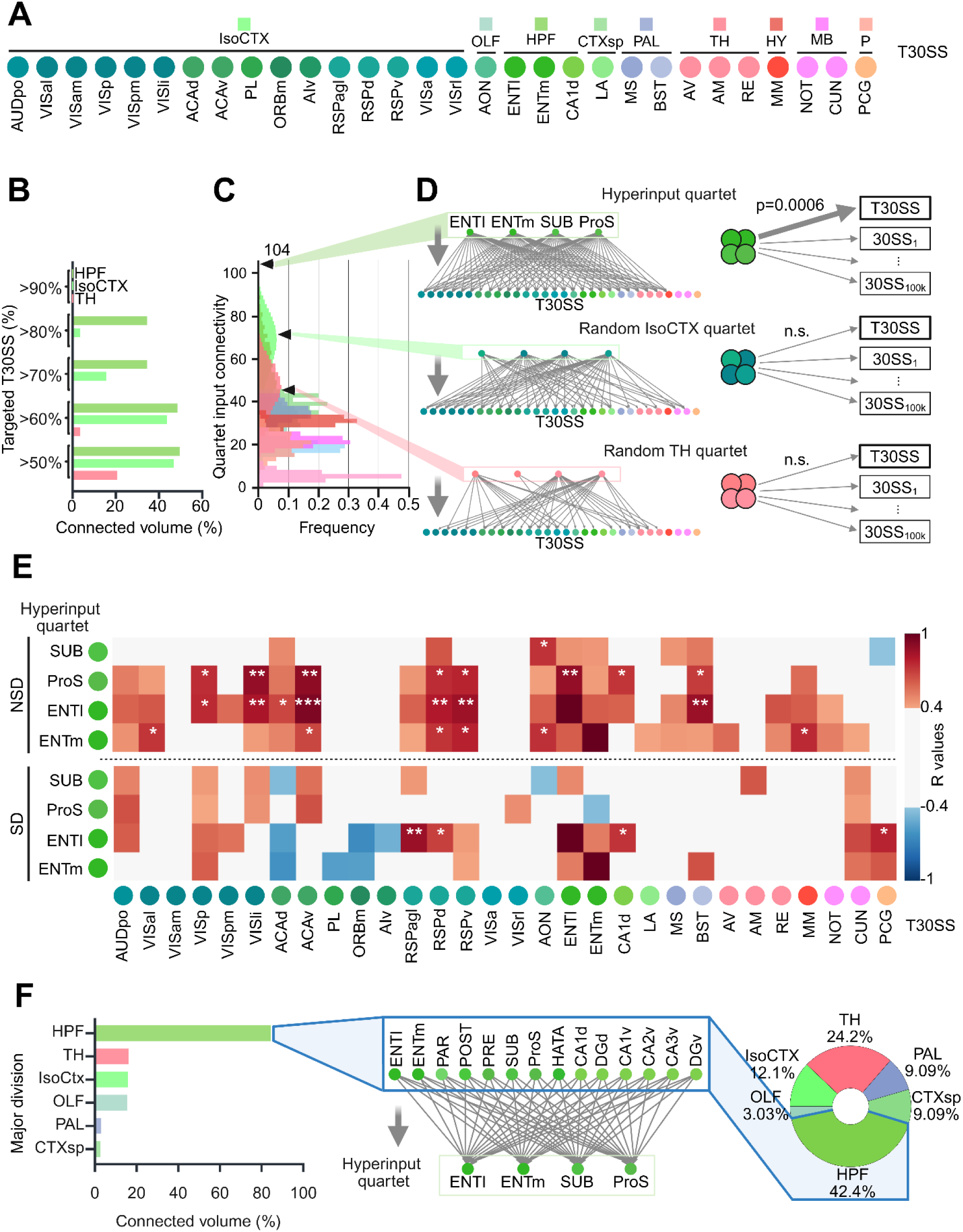
The hippocampal formation is an input hub targeting the brain network impaired by SD . **(A)** List of the thirty brain regions (T30SS) showing the most prominent cFos differences in SD vs NSD mice. Colors reflect anatomical organization by major division **(B)** Identification of hyperinputs to the T30SS. Percentage of T30SS receiving projections from each summary structure, stratified by the corresponding major division volume. Only divisions for which at least one summary structure is connected to >50% of T30SS are shown; all other major divisions exhibit <50% and are not shown. The HPF shows the largest volume highly connected to T30SS, followed by IsoCTX and TH. **(C)** Distribution of input connectivity to T30SS for each quartet (combination of 4 summary structures stratified by major division). Quartet input connectivity (y axis) is calculated as the sum of the number of T30SS targeted by each component of the quartet. The “hyperinput quartet” (ProS, SUB, ENTl, and ENTm) show the highest connectivity to the T30SS accounting for 104 of 120 total possible projections to T30SS. Colors indicate major divisions as depicted in A. **(D)** Left: Diagram of the hyperinput connectivity to the T30SS (Top), and of two additional quartets randomly drawn within Isocortex and TH (bottom). Right: comparison of the hyperinput quartet→T30SS connectivity versus its connectivity to random 30-region sets (top, p≤0.00006). Randomly selected quartets from cortical or thalamic regions do not show such specificity (IsoCTX p=0.30964; TH p=0.82876). **(E)** Functional connectivity analysis between the hyperinput quartet (rows) and the T30SS (columns) in NSD and SD mice following memory recall. Pearson correlation matrices showing inter-regional correlations of cFos density. Colors reflect Pearson correlation coefficients and starts indicate correlation p values. *p < 0.05; **p < 0.01; ***p < 0.001; ****p < 0.0001. **(F)** Input connectivity to the hyperinput quartet. Left: bar plot showing the percentage volume of each major brain division connected to all four hyperinput quartet regions. Middle: focus on the hippocampal formation, highlighting the subregions connected to all four regions of the hyperinput quartet. Right: major division of origin of the shared projections to the hyperinput quartet. Notably, 42% of structural inputs come from the HPF. Detailed description of statistical analysis can be found in the Supplementary Table 2. Created using BraiAn, BrainGlobe, and BioRender.com.

### Object-location memory task

To reduce stress and acclimate animals to experimental procedures, mice were handled for two days (4 min/day, by two experimenters). On the second day, mice were weighed and given a mock intraperitoneal (i.p.) injection (one needle insertion with an empty syringe) if the subsequent experiment required an injection (Fig. 3, S7, 4). The next day, they were habituated for 5 min to the empty OLM arena. The OLM task took place in a PVC arena (40 × 30 × 30 cm) with two large spatial cues on opposing short walls (checkerboard or striped patterns)^8,28,43^. During habituation, no objects were present. In the training session, two identical objects were placed symmetrically ∼7.5 cm from one of the long and one of the short side walls, and mice explored for 10 min after being introduced from the same starting position. After training, mice returned to their home cage and either left undisturbed (non–sleep deprived, NSD) or kept awake for 6 h in the home cage (sleep deprived, SD). In the recall session on the following day, one object was displaced ∼15 cm along a straight line, and mice were allowed to explore the arena and objects freely for 10 min. Different sets of objects were used for different animals, but the different sets were similarly distributed across treatment groups. The original location of the objects as well as the choice for the displaced object (left or right/up or down) was randomized across animals and treatment groups. The arena and objects were cleaned with 70% ethanol between trials. Habituation, training and test sessions took place at the beginning in the first two hours of the light phase. Memory performance during the test session was quantified using the d2 discrimination index: (time exploring relocated object – time exploring non-relocated object) / total objects exploration time. Exploration was scored using BORIS software^44^, with exploration defined as nose contact or orientation within 1 cm to the object. Mice that climbed onto objects with all four paws or escaped the arena were excluded from analysis.

**Figure 3.**
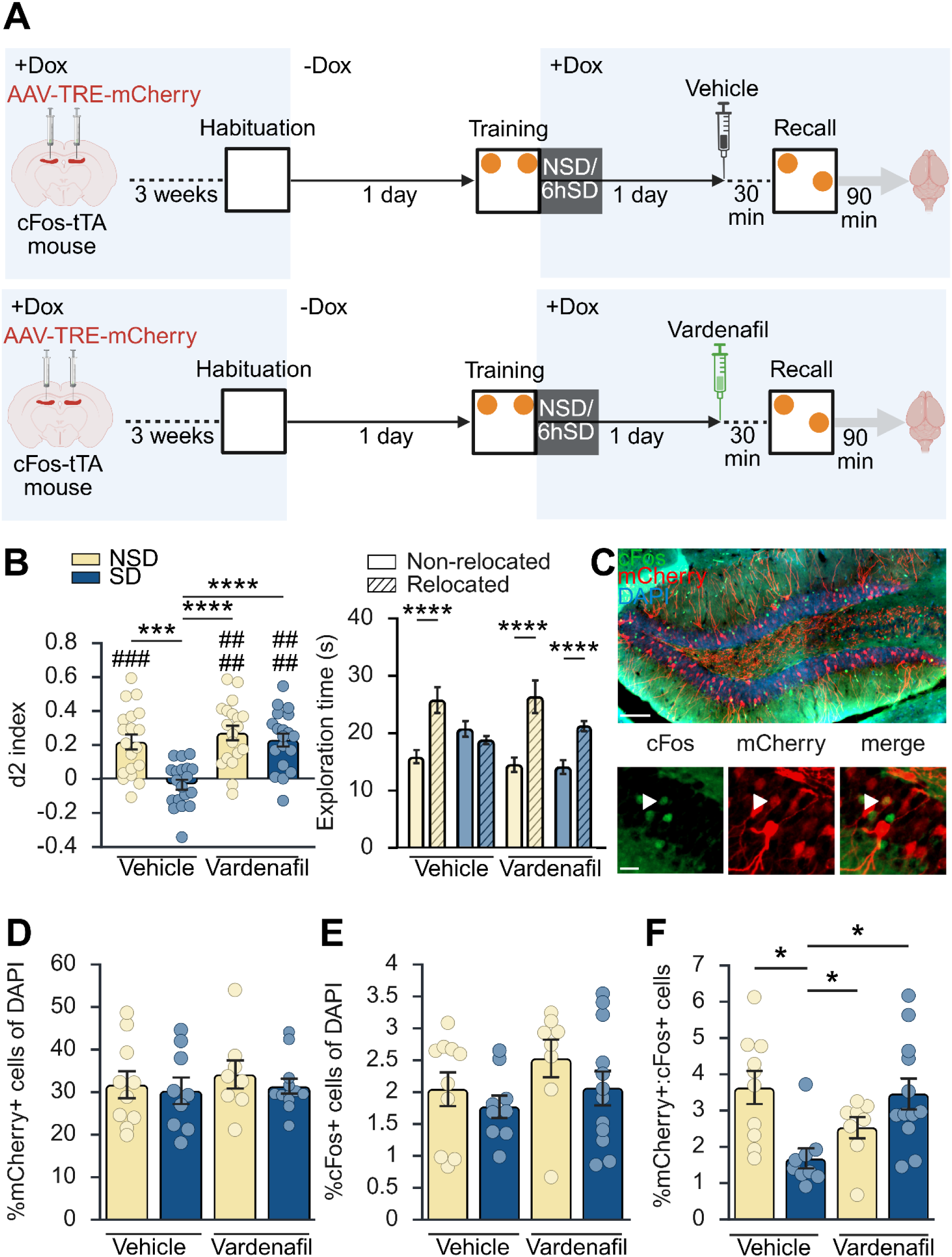
Vardenafil treatment rescues object-location memory deficits caused by sleep deprivation. **(A)** Diagram to explain the experimental procedure, n=19. **(B)** OLM performance. Left: discrimination (d2) index. Only the combined treatment group performed above chance during the test session. Right: During testing, NSD controls and SD-vardenafil mice preferred the relocated object, whereas SD-vehicle mice failed to discriminate. Raw values are presented for ease of interpretation; however, statistical analysis for the right panel was performed on log₁₀-transformed data to satisfy normality assumptions. **(C)** DG engram cell analysis. Representative coronal sections showing mCherry+ (red), cFos+ (green), and colocalized cells (yellow; scale bar:50 µm; C). **(D-E)** Total percentages of mCherry+ and cFos+ cells did not differ across groups. **(F)** Colocalization of mCherry+ and cFos+ cells was reduced in SD-vehicle mice, consistent with impaired engram reactivation. Data are presented as mean ± SEM. Statistical significance is denoted as */#p<0.05, **/##p<0.01, ***/###p<0.001, ****/####p<0.0001. # indicates significantly different from zero. Created using BioRender.com.

**Figure 4.**
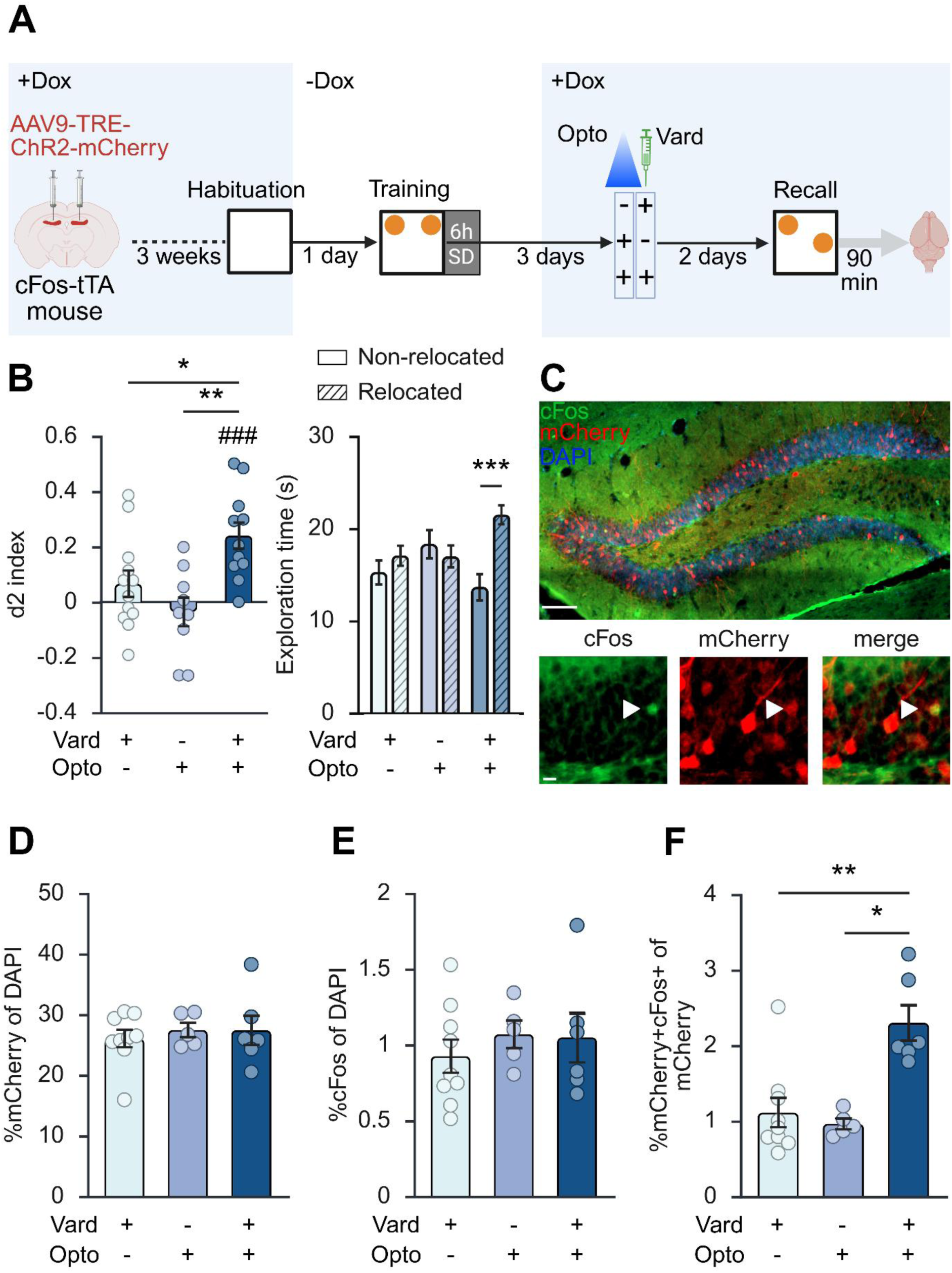
Combined optogenetic and vardenafil treatment leads to permanent memory recovery by increasing engram reactivation. **(A)** Schematic representation of the experimental timeline. **(B)** Left: discrimination (d2) index (n = 9–12 per group). Only the combined treatment group performed above chance during the test session. Right: Exploration time of the two objects during recall. Only the combined treatment group preferentially explored the relocated object. Raw values are presented for ease of interpretation; however, statistical analyses for the right panel were performed on log₁₀-transformed data to satisfy normality assumptions. **(C)** Representative coronal brain section of the DG showing mCherry (red signal; cells active during training), cFos (green signal; cells active during recall), and overlapping mCherry+cFos+ (yellow signal) in brain tissue of mice subjected to the experiment. **(D-E)** Quantification of mCherry⁺ (F) and cFos⁺ (G) cells over the total number of DAPI cells, showing no differences among groups (n=5-9 per group). **(F)** Overlap of mCherry⁺/cFos⁺ cells, with significantly higher co-localization in the combined treatment group. Data are presented as mean ± SEM. *P≤0.05, **P≤0.01, ***P≤0.001, ****P≤0.0001. # indicates significantly different from chance level. Created using BioRender.com.

### Sleep deprivation

Animals were subjected to a single 6-hour SD session between the first few hours of the light phase, starting immediately after the training trial in the OLM task. After this, mice were left undisturbed in the home cage for 18 hours until the testing session. SD was achieved by using the gentle stimulation method, which involves gently tapping on the cage or shaking the cage, and is considered only mildly stressful, as it does not increase glucocorticoid levels beyond baseline levels observed across the circadian cycle^32,45–49^. This SD method has been validated through EEG recordings, which show a complete loss of REM sleep and approximately 95% of NREM sleep^50^.

### Brain extraction and fixation

Ninety minutes following the behavioral testing (Fig. 1, 3, 4), mice were transcardially perfused with 0.9% NaCl containing heparin, followed by 4% paraformaldehyde in 0.01M phosphate-buffered saline (PBS). Brains were post-fixed in PFA for 48 hours at 4°C. Subsequently, brains were washed with 0.01M PB for 24 hours and then transferred to a 30% sucrose solution for cryoprotection for 48 hours, with the solution renewed daily. Finally, brains were frozen using liquid nitrogen and stored at -80°C.

### Whole-brain immunofluorescence staining

For whole-brain cFos immunostainings (Fig. 1), 40 micron coronal brain sections were cut with a sliding cryostat. Sections were stored in 48-well plates free-floating in 1X PBS with 1% sodium azide at +4°C. Immunofluorescence staining was performed on free-floating sections (1 in 5 whole-brain series). First, sections were washed 2 times in 1X PBS at room temperature (RT, 5 minutes each) and subsequently incubated in blocking solution at RT (1X PBS, Triton 0.3% and bovine serum albumin, 1%) for 90 minutes under constant shaking. Next, the sections were incubated with primary antibody (rabbit anti-cFos, 1:5000, Synaptic Systems, #226 008) in 1X PBS with 1% BSA and Triton 0.1% for 48 h at 4°C under constant shaking. After incubation, sections were left at RT for 15 minutes, thoroughly washed in PBST (Triton 0.1%) and subsequently incubated with Alexa-conjugated secondary antibody: Donkey anti-rabbit 647 (1:1000, Invitrogen, A31573) diluted in PBST (Triton 0.1%) at RT for 2 hours under constant shaking. Sections were then washed 3 times (5 minutes each) with 1X PBS and then mounted on superfrost glass slides (ThermoScientific) with DAPI Vectashield (Vector Labs, H-1200-10). Images were acquired with a Zeiss Axioscan microscope Z1 equipped with a Hamamatsu Orca Flash 4 camera (2048×2048 pixels, 6.5 µm pixel size) with a 20×/0.8 Plan Apochromat objective. The resulting image’s pixel size is 0.325 µm.

### Whole-brain image analysis

Whole-brain datasets were aligned to the 10µm Allen Brain Atlas CCFv3^51^, in which we added a division between the dorsal and ventral hippocampus. Atlas registration was performed using Aligning Big Brains and Atlases (ABBA)^14,52–55^. Slice positioning on the anterior-posterior axis and angle correction were obtained by DeepSlice^53^ and subsequently checked and manually adjusted when needed. For all sections, atlas registrations were first performed automatically by affine and spline registrations^54,55^, and subsequently manually refined using BigWarp^52^ to maximize precision. At least two independent experimenters checked atlas registration accuracy. All atlas annotations were imported into QuPath (v0.5.1;^56^). Quantification of cFos+ cells was performed on 16-bit grey scale images, using BraiAn extension for QuPath^14^. Positive cell detection, segmentation, and classification were performed following the procedures described in Chiaruttini et al. 2025^14^. Damaged or misaligned tissue portions were excluded from further analysis.

### Whole-brain data analysis

Statistical analyses and whole-brain visualization were performed in Python using BraiAn^14^. First, we set atlas parcellation granularity at “summary structures” as defined by Wang et al. 2020^51^ (whole brain atlas parcellated in 316 regions) and performed a quality control screening across the whole dataset by computing the coefficient of variation of cFos+ cell density for each summary structure across different sections of the same animal. When the variation of cFos positive cell density changed drastically between sections from the same animal, images were manually checked for potential artifacts. Subsequently, we calculated the number of positive cells for each summary structure by summing cFos+ cells across all sections. The same was done with the associated structures’ area in mm². All subsequent analysis was based on per region density measure (cFos+/mm2). All regions missing for at least one subject, as well as the cerebellum, fiber tracts and the ventricular system were excluded from the analysis. For assessing global differences between groups we used Mean-centered Task Partial Least Squares (PLS) analysis^57^ as implemented in BraiAn^14^, in which statistically significant differences are assessed through permutation test, while the reliability of the contribution of each nonzero region salience is determined using bootstrap estimates of the salience standard errors^58^. Brain division into four functional circuits was obtained on the basis of existing literature: memory-associative^59–66^; emotional-motivational-arousal^67,68^; stimulus processing^69–72^; sensory motor^73–80^. Brain areas assigned to each functional circuit are listed in Supplementary Table S1. Global differences between SD and NSD mice for all the summary structures as well as for each functional network were independently assessed by PLS, through 10K permutations. The reliability of the scores of each region in the memory-associative network was performed by bootstrap, through 10K resamples. The resulting estimates of the regions’ salience scores were interpreted as Z-scores, with Z ≈ ±1.96 taken as statistically stable (p < 0.05, two-tailed).

For network connectivity analysis, we first used the salience defined by PLS calculated over the summary structures, stabilized through 10K bootstrap resamples, and identified the 30 most prominent regional contributors to the whole-brain contrast between SD and NSD mice. These Top 30 Summary Structures (T30SS) were identified by ranking all summary structures (n=214; removing those with missing cFos quantifications) according to the highest absolute magnitude of their bootstrap-derived Z-scores. For connectome data analysis, we utilized a high-resolution data-driven model of the mouse connectome, derived from 428 anterograde tracing experiments in young adult wild-type mice^81^. To build an unweighted network, we used the “normalized connectivity density” c_i,j_ defined by the model for any couple of brain structures <i,j>. Note that c_i,j_≠c_i,j_. This index defines the level of anatomical connectivity, and for its binarization we only kept the connections <i,j> such that log(c_i,j_) > -4.5. Connected volume was computed by measuring the summary structures’ outdegree to the T30SS, in relation with the corresponding volume of the structure within its major division. The volume of each structure and major division was defined by the number of voxels of the corresponding annotation within the 3D atlas. Input connectivity to a set *B* of regions for another set *A* of regions was defined as the sum of each structure’s outdegree to the 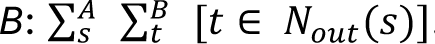. To test the specificity of an hyperconnection of a set *A* of regions to T30SS, we calculated the input connectivity for *A* to 100,000 random sets of 30 summary structures (30SSi) and compared them to the input connectivity to the T30SS. A bootstrap p-value was calculated as the fraction of these 100,000 random sets that exhibited a higher input connectivity. Importantly, for each 30SSi we maintained the same anatomical distribution of the T30SS (Fig. 2A; 16 IsoCTX, 1 OLF, 3 HPF, 1 CTXsp, 2 PAL, 3 TH, 1 HY, 2 MB, 1 P). Cross correlations were performed with the Pearson method on the cFos density in every structure of each group, following a normalization on the cFos density of the whole brain. Regions for which the quantification was present in less than 4 animals were discarded from the cross correlation. For each correlation of two structures, we also performed a test of the null hypothesis that the distributions underlying the samples are uncorrelated and normally distributed.

### Virus construct and packaging

To label memory engram cells (Fig. 3 and Fig. 4), we bilaterally injected an adeno-associated virus (AAV9-TRE-ChR2-mCherry, titer: 3.75e14, 200 nL; Penn Vector Core, Philadelphia, US) in the dentate gyrus (DG) of c-fos-tTA mice using the following coordinates: -2.0 mm anterior-posterior (AP), 1.3 mm ± medial-lateral (ML), -1.8 mm dorsal-ventral (DV). Channelrhodopsin-2 (ChR2) and mCherry expression were controlled by the tetracycline response element (TRE), as previously described^28^. In this way, cFos-driven expression of tTA induces ChR2 and mCherry expression, which can be suppressed by keeping animals on a Dox-containing chow (*i.e.*, cFos expression triggers ChR2 and mCherry expression only in the absence of Dox).

### Viral injection and optic fiber implants surgery

All surgeries were performed in a stereotaxic frame (Kopf Instruments) following established procedures^28^. Mice were anesthetized with isoflurane (5% induction; 1.8–2.0% maintenance) and placed on a heating pad to maintain body temperature. Carprofen (0.1 mL/10 g at 0.5mg/ml concentration) and lidocaine (0.1 mL at 20mg/ml concentration) were administered immediately after induction. Small bilateral holes were drilled in the skull using a 0.5 mm diameter microdrill at the appropriate locations, and viral solutions were injected using a 10 μL Hamilton syringe with a 33G nanofil needle (WPI). The needle was lowered to the pocket site at –1.9 mm (1 min), raised slightly to –1.8 mm (1 min), and the virus was delivered at 70 nL/min. After injection, the needle was held for 1 min at –1.8 mm and –1.7 mm before withdrawal. For experiments involving only viral labeling of engram cells (Fig. 3), the skin was closed with sutures. For optogenetic experiments (Fig. 4), optical fibers were implanted at –2.0 mm AP, ±0.13 mm ML (fiber length 0.175 mm) and secured with screws and C&B Metabond. At the end of surgery, mice received 0.1 mL saline and were allowed to recover for 3 weeks before starting the behavioral experiments.

### Optogenetic stimulation protocol

For the experiment involving optogenetic stimulation (Fig. 4), DG engram cells were optogenetically stimulated with 15 ms pulses at 20 Hz for 3 minutes. Stimulation was induced using a 473nm laser (CrystaLaser) driven by a TTL input (Doric lenses). The power delivered at the DG was 10-15 mW. All optogenetic laser stimulation took place in the home cage of the test mouse, which was connected to the laser cables for 5 minutes.

### Drug preparation

Drug preparation was performed as previously described^42^. Vardenafil hydrochloride powder was dissolved in the vehicle solution (methyl cellulose tylose solution (0.5%) and 2% Tween80) no longer than 24 hours before the injection. All components were obtained from Sigma Aldrich, Zwijndrecht, the Netherlands. The injection volume was calculated in proportion to the body weight, 2 ml/kg. Vardenafil was administered i.p. at a dose of 0.3 mg/kg. Dose and injection volume were based on the previous work with vardenafil in memory studies^42^. Injections were given 30 minutes before the OLM test in the studies without optogenetic stimulation (Fig. 3 and Fig. S7). In the study combining optogenetic stimulation and pharmacological treatment, the injection was given immediately after the optogenetic stimulation (Fig. 4). At this time point PDE5 inhibition was shown to be most effective for memory enhancement in mice^37,40^.

### Engram cells immunofluorescence staining

For pharmacological and optogenetic experiments (Fig. 3, 4), coronal brain sections were cut at a thickness of 20 microns with a sliding cryostat. Immunofluorescence staining was performed to visualize cFos and mCherry expression in hippocampal tissue sections. Cups containing 3 to 4 dorsal brain sections were washed three times with 0.01 M PBS for 5 min each time and incubated for 20 min in 0.3% H202 in PBS on a shaker at RT. Then, sections were washed 3 times x 10 min with PBS and incubated with 5% normal donkey serum (NDS) in PBST (0.2% TritonX100) for one hour (to block nonspecific binding). Next, the sections were incubated with primary antibodies—mouse anti-cFos (1:1000, Abcam, #ab208942) and rabbit anti-mCherry (1:1000, Invitrogen, #PA5-34974)—in PBS with 1% NDS and 0.2% Triton-X100 for 3 nights at 4°C. After primary antibody incubation, all the sections were washed 2 times for 5 min and 2 times for 10 min with PBS at RT and then incubated for four hours (at RT and protected from light) with secondary antibodies—donkey anti-mouse AF488 (1:400, Jackson Labs, #A-21202) and donkey anti-rabbit AF555 (1:500, Jackson Labs, #A-31572)— diluted in 1% NDS. Sections were then washed 4 times for 5 minutes each with PBS, mounted on glass slides, and covered with DAPI Vectashield (Vector Labs, H-1200-10). Fluorescent signals were visualized using a Zeiss AXIO Observer microscope with 20x magnification. Images were captured using DAPI, mCherry, and cFos filters with exposures set to avoid overexposure while maintaining clear visibility of labeled cells. Images were analyzed using Zeiss software to stitch and process images.

### Engram cell counting

The number of mCherry- and cFos-positive cells was assessed in 3 coronal sections per animal to quantify their expression and colocalization in the dentate gyrus (DG; Fig. 3, 4). Fluorescent images were obtained using a Zeiss AXIO Observer microscope and analyzed in QuPath (v.0.4.3). A pre-developed script (StarDist) has been optimized to automate cell counting^82^. The cell body layer within the dorsal DG region was defined as the region of interest (ROI). The StarDist script was then adapted to use DAPI staining to identify the total number of nuclei. In addition, a classifier was trained in QuPath to identify and discriminate cFos and mCherry positive cells and applied to all sections. Colocalization was defined as cells positive for both cFos and mCherry. The following formulas were used: ((cFos+ cells + cFos+mCherry+ cells) / DAPI cells)*100; ((mCherry+ cells + cFos+mCherry+ cells) / DAPI cells)*100; and (cFos+mCherry+ cells / mCherry+ cells)*100. Manual verification of cell counts was performed in a blinded manner. Damaged brains or slices, or those with mCherry expression restricted to one hemisphere, were excluded. All analyses were run using the same parameter-configuration file to ensure full reproducibility across animals and groups.

### Statistical Analysis

Data were analyzed by unpaired t-tests, one-way and two-way ANOVAs, and post-hoc tests as appropriate and indicated for each experiment. All data are represented as mean ± SEM using JASP software (Version 0.19.3.0; https://jasp-stats.org/) and Prism 10 (GraphPad Software, La Jolla, CA). Graphical illustrations were generated using BioRender (BioRender, 2025; https://biorender.com/). The statistical significance was set at p < 0.05. Detailed statistical tests and data with exact p values are listed in supplementary Table S2.

## Results

### Sleep deprivation-induced impairment of OLM recall is associated with decreased activation of memory-related brain-wide networks

We hypothesized that SD following memory encoding causes impaired memory retrievability, associated with a dysregulation of large-scale network activity, extending beyond localized hippocampal deficits. To test this hypothesis, C57BL/6J male mice were subjected to a single SD period of 6 hours immediately after the training trial in the OLM task, using the gentle handling method, considered only mildly stressful^32,45–49^. After this period, mice were left undisturbed in the home cage for 18 h until the test session for memory recall (24 hours after training; Fig. 1A). Consistent with previous findings^8,28,43^, during recall, NSD controls exhibited a significant preference for the relocated object, indicative of an intact spatial memory, whereas SD mice failed to discriminate between objects, reflecting impaired memory retrieval (Fig. 1B, left: unpaired t-test: SD vs. NSD t22 = 2.291, p = 0.0319; right: two-way RM ANOVA, F1,22 = 5.283, p = 0.0314). During training, all groups displayed comparable exploration of both objects, indicating no innate preference (Fig. S1; N.S.).

To investigate how such impaired memory retrieval is reflected in network activity at the systems level, we performed brain-wide activity mapping from brains collected 90 minutes following OLM recall in SD and NSD animals by measuring the expression of the immediate early gene cFos (Fig. 1C, D; Chiaruttini et al.^14^). Interestingly, the global density of cFos-positive cells did not differ between groups, indicating an absence of global activity changes (Fig. 1E, S2, S3, S4). To test whether SD-related effects may emerge within specific brain networks rather than at the whole-brain scale, we defined four macro-circuits associated with distinct functional processes based on the existing literature^59,64,67,71,75,77^, namely: stimulus processing, motivational-emotional-arousal, sensory-motor, and memory-associative circuits (a detailed description of the regions composing each circuit is available in Supplementary Table 1). PLS analysis comparing cFos density in the two groups revealed that, among the four networks, only the memory-associative network showed significant differential activity in SD vs NSD mice (Fig. 1F; mean-centered task PLS, p = 0.0174, 10 000 permutations). Within this network, a number of cortical, hippocampal, and subcortical regions showed decreased cFos in SD compared to NSD mice (Fig. 1G). For instance, SD mice had reduced cFos density in the hippocampal CA1 as well as the retrosplenial, anterior cingulate, and entorhinal cortices: regions critically implicated in object and spatial representations^83–87^.

Because our whole-brain mapping analyses of impaired spatial memory recall revealed a network-specific activity impairment localized to memory-relevant circuits, we next investigated the functional connectivity organization of this network. For this, we combined our whole-brain cFos mapping results (Fig.1) with structural connectivity data from the Allen Institute mouse connectome atlas^81^ to identify putative hub regions that may serve as central upstream orchestrators of the distributed memory network impaired by SD.

The SD impaired network was defined as the pool of thirty regions (Top 30 Summary Structures, T30SS) showing the largest differences between SD and NSD mice across the whole brain (Fig. 2A; obtained by whole-brain PLS analysis; for analytical details see Materials and Methods) and included mostly cortical and subcortical regions within the memory-associative macrocircuit (Fig. 1F-G, 2A). To determine whether we could identify candidate input hubs extensively and selectively targeting the T30SS (hereafter referred to as “hyperinputs”), we assessed the brain-wide monosynaptic inputs to all the 30 brain structures composing the T30SS and looked for regions targeting the highest number of the T30SS and the lowest number of non-T30SS regions. We ranked all T30SS inputs by the number of targeted T30SS (Fig. S5), grouped them by major brain division and assessed which major division contained the highest proportion of inputs hubs (defined as regions targeting at least 26 out of the thirty T30SS. Fig. 2B). This analysis identified the hippocampal formation (HPF) as the major division most prominently targeting the T30SS. Specifically, four HPF subregions, representing 30% of the total HPF volume, projected to at least 26 of the 30 T30SS regions (86.5%; Fig. 2B). The HPF was followed by the isocortex (IsoCTX), which showed 0.6% of its volume connected to 27 T30SS regions and by the thalamus (TH), with 3.7% of its volume connected to 20 T30SS regions. All other major divisions projected to less than 50% of the T30SS. Therefore, the HPF emerged as the major division with the largest proportion of its volume targeting the majority of T30SS regions, suggesting a potentially central role in coordinating activity across these structures.

To verify the unique structural connection between the four HPF subregions and the T30SS, we quantified the connectivity distribution to the T30SS across all combinations of four summary structures (quartets; Fig. 2C, stratified by major divisions). This analysis identified four hippocampal regions (the presubiculum (ProS), subiculum (SUB), lateral entorhinal cortex (ENTl), and medial entorhinal cortex, (ENTm)) that collectively projected to nearly all T30SS regions, except for PCN, CUN, and NOT located in the midbrain and pons (Fig. 2D, left). We hereafter refer to these four hippocampal regions as the “hyperinput quartet”. Notably, this hyperinput quartet exhibited the highest connectivity to the T30SS compared with all other possible four-region combinations within each major division (Fig. 2C).

We subsequently raised the question whether the identified hyperinput quartet is specifically hyperconnected to the T30SS and not unspecifically hyperconnected to the whole brain. To test this hypothesis, we calculated the input connectivity of the hyperinput quartet to 100,000 random sets of 30 summary structures (30SS_i_) and compared this to the connectivity to the T30SS. The analysis confirmed that the input connectivity of the hyperinput quartet was specific to the T30SS (p=0.00006, Fig. 2D right). When each region of the hyperinput quartet was examined individually, all remained selectively connected to the T30SS (SUB: p=0.00004; ProS: p=0.00017; ENTl: p=0.01359; ENTm: p=0.02326). Remarkably, this hyperconnectivity to the T30SS did not emerge in other randomly generated input quartets from other major divisions (random cortical quartet, p=0.30964; random thalamic quartet, p=0.82876; Fig. 2D, right), thus highlighting the unique identity of the hyperinput quartet as input hub to the T30SS. Collectively, these findings identify an input hub of four regions that may globally influence the specific network associated with impaired retrievability of OLMs formed under sleep deprivation. After having identified the input hub by its unique anatomical connectivity to the T30SSt, we investigated whether its functional connectivity to the T30SS was affected by SD by computing pairwise inter-regional covariance of cFos density between the four hyperinput regions and the T30SS in the two groups. Notably, in NSD mice, cFos activity in the hyperinput quartet showed significant covariation with most T30SS, while such a correlation pattern was lost in SD mice (Fig. 2E). These results suggest that SD during consolidation may weaken the synchronized activity between the hyperinput quartet and its downstream targets at the time of memory recall. Collectively, whole-brain cFos mapping revealed that the memory retrievability impairment induced by SD during the initial phases of memory consolidation, is associated with reduced cFos expression within memory-associative circuitry and identify the HPF as a possible orchestrator of such network changes. Indeed, anatomical connectivity analysis identified four hippocampal regions (hyperinput quartet) as an input hub to the SD-impaired memory network (T30SS). Interestingly, this identified input hub comprises four regions that together form the principal output system of the hippocampal formation^88–92^ with 84.7 % of its volume sending projections to all four regions of the hyperinput quartet (Fig. 2F; 14 out of 19 HPF regions; supplementary Table S3) and representing 42% of its brain-wide input connectome.

### Sleep deprivation-induced memory recall deficits are associated with reduced DG engram reactivation and vardenafil administration upon memory recall rescues the associated memory deficits and enhances DG engram cells reactivation

In the previous section, we showed that the majority of the input regions projecting to the hyperinput quartet were confined to hippocampal formation (Fig. 2F). Several of these regions, despite unaltered cFos expression between groups (Fig. 1G, S2) are well established as critical inputs for memory recall. Although additional regions may warrant investigation, the dentate gyrus (DG) is of particular interest because it provides input to all four hyperinput regions. Moreover, prior literature identifies the DG as the principal gateway to the trisynaptic hippocampal circuit and a key regulator of hippocampus-dependent memory retrieval^93^. Consistent with this role, studies in other models of amnesia have shown that retrieval deficits have been associated with impaired reactivation of DG engram cells^94–97^. We therefore examined whether SD specifically disrupts DG engram reactivation, potentially accounting for the observed retrieval impairment. To address this question, cFos tTA mice underwent OLM training, allowing us to tag DG engram cells activated during learning, followed by 6 hours of SD. The day after, mice underwent recall and brains were collected 90 minutes after (Fig. 3A, n=19). We then quantified neurons expressing mCherry (cells active during training), and cFos (cells active during recall). Colocalization of mCherry+ and cFos+ cells indicated re-engagement of the original memory engram cells during recall, with higher percentages reflecting successful memory retrieval^95–98^. SD mice that failed to properly recall the OLM (Fig. 3B), showed no differences in the percentage of cFos+ and mCherry+ cells between groups (Fig 3C-E), confirming prior results that overall cFos levels did not differ in the DG between groups. Interestingly, we found decreased colocalization of mCherry+ and cFos+ cells in the DG of SD mice (Fig. 3F), indicating that the negative effects of SD on memory recall are reflected in a selective failure to reactivate memory engram cells.

Based on these findings, we next sought to determine whether enhancing cGMP–PKG signaling via the FDA-approved PDE5 inhibitor vardenafil could rescue OLM performance, and if so, whether this would happen by promoting reactivation of these memory engram populations. To address this, cFos tTA mice underwent OLM training followed by 6 hours of SD. Thirty minutes before the retention test on the following day, animals received an i.p. injection of vardenafil, and brains were collected 90 minutes after recall (Fig. 3A). During training, all groups displayed comparable exploration of both objects, indicating no innate preference (Fig. S6B; N.S.). During testing, d2 index measurements showed that vardenafil could restore memory performance in SD mice (Fig. 3B left: two-way ANOVA, interaction: F_1,72_ = 6.917, p = 0.0104; post-hoc Bonferroni: NSD-vehicle vs. SD-vehicle p = 0.0002, NSD-vardenafil vs. SD-vehicle p < 0.0001, SD-vardenafil vs. SD-vehicle p < 0.0001). Consistently, the exploration times of the two objects during testing confirmed that NSD controls and SD-vardenafil mice showed a clear preference for the relocated object, consistent with intact spatial memory, whereas SD mice failed to discriminate between objects, indicating impaired memory retrieval (Fig. 3B right: two-way RM ANOVA, interaction: F_3,72_ = 11.34, p < 0.0001; post-hoc Bonferroni: NSD-vehicle p < 0.0001, NSD-vardenafil p < 0.0001, SD-vardenafil p < 0.0001, SD-vehicle N.S.). PDE5 inhibition did not improve cognitive performance in non-SD conditions, likely because baseline cyclic nucleotide levels are sufficient for normal hippocampal function. Similar results were observed in C57Bl/6J mice (Fig. S7), highlighting the reproducibility of the data and the efficacy of vardenafil across strains. To determine whether the beneficial effect of vardenafil treatment on memory reflects a selective re-engagement of memory engram cells during recall, we quantified neurons expressing mCherry, cFos and their colocalization (Fig. 3C). The total numbers of mCherry+ or cFos+ cells did not differ between groups (Figs. 3D-E; two-way ANOVA, interaction: N.S.; n = 9–12). By contrast, SD mice treated with vardenafil showed significantly higher colocalization with respect to mice that were SD with impaired OLM recall (Fig. 3F; two-way ANOVA, interaction: F1.35 = 4.458, p = 0.0420; post-hoc Bonferroni: SD-vehicle vs. NSD-vehicle p = 0.0119, SD-vehicle vs. NSD-vardenafil p = 0.0179, SD-vehicle vs. SD-vardenafil p = 0.0119; n = 9–12).

Taken together, these findings suggest that SD disrupts memory accessibility by reducing engram cell reactivation, rather than altering overall cFos activity in the DG, thereby impairing the retrieval of previously encoded memories. Moreover, these data also suggest that vardenafil has beneficial effects on memory retrieval following SD. These effects are not driven by a global increase in neuronal activity within the DG, but rather by the selective reactivation of the specific neuronal ensemble necessary for appropriate memory recall to occur.

### Optogenetic DG engram stimulation followed by vardenafil treatment leads to persistent memory recovery by increasing engram reactivation

In the present study, successful memory retrieval in sleep-deprived mice was observed when vardenafil treatment immediately preceded the retrieval test. We therefore aimed to persistently restore long-term access to OLM, enabling successful natural recall days after intervention. Akkerman et al.^39^, established that transient short-term memories in rodents can be converted into more robust memories through systemic treatment with PDE5 inhibitors within 0-45 minutes following training, corresponding to the early consolidation window^39^. We therefore combined optogenetic reactivation of the OLM engram formed under SD with subsequent systemic administration of the PDE5 inhibitor vardenafil to determine whether this approach could enable lasting retrieval of the memory and promote a more natural recall (*i.e.*, without the need for additional optogenetic stimulation or drug treatment prior to testing days later). To address this, mice were subjected to 6 hours of SD following training. Three days later, animals were subjected to one of the three experimental conditions: (1) laser ON followed by i.p. vehicle injection (ON_veh), (2) laser OFF followed by vardenafil i.p. injection (OFF_Vard), or (3) laser ON followed by vardenafil i.p. injection (ON_Vard). Two days after treatment, mice underwent the test trial (Fig.4A). In our study, the “learning episode” is replaced by optogenetic reactivation of the engram, which is then reconsolidated in the presence of boosted cGMP signaling. During the training trial, all mice explored both objects equally, showing no preference for either object (Fig. S6C; Two-way RM ANOVA, N.S.). Mice that received solely optogenetic stimulation or vardenafil treatment failed to detect the spatial novelty indicating that they failed to recall the previously acquired spatial memories. In contrast, mice receiving the combined treatment performed above chance level during testing suggesting that they successfully detected the spatial novelty (Fig. 3D; One-way ANOVA, F_2,29_ = 7.383, p = 0.0026; one-sample t-test vs. chance: ON_vard, t_10_ = 4.966, p = 0.0006; OFF_vard and ON_veh: N.S.). Post-hoc comparisons further demonstrated that ON_vard mice exhibited significantly higher discrimination index values than both OFF_vard (p = 0.0328) and ON_veh conditions (p = 0.0016). Analysis of exploration time of objects corroborated these findings: mice that received only one of the two treatments explored the relocated and non-relocated objects equally (Fig. 3B left: OFF_vard and ON_veh: N.S.). A significant interaction between condition and target emerged (Two-way RM ANOVA, F_2,29_ = 7.303, p = 0.0027), and post hoc analyses indicated that the combined treatment group exhibited significantly greater exploration of the relocated object relative to the non-relocated one (ON_vard, p = 0.0001). Consistent with our expectations, vardenafil treatment alone failed to produce a lasting recovery of memory, as cGMP elevation in the absence of memory-related neural activation is insufficient to drive reconsolidation processes^37^.

To determine whether restored OLM was associated with altered engram reactivation in the DG, we quantified mCherry⁺, cFos⁺, and double-labeled mCherry⁺:cFos⁺ neurons (Fig. 4C). The numbers of mCherry⁺ and cFos⁺ cells did not differ across conditions (Fig. 4D-E; N.S.). In contrast, engram reactivation, indexed by mCherry⁺cFos⁺ colabeling, was significantly increased in mice that received the combined treatment (Fig. 4F; Kruskal–Wallis, H_2_ = 9.833, p = 0.0029), with post-hoc test indicating greater overlap compared to both OFF_vard (p = 0.0099) and ON_veh (p = 0.0410). Together, these findings indicate that the combined approach enables a persistent recovery of access to memories formed under SD, facilitated by increased reactivation of DG engram cells in the absence of global changes in cFos activity. This not only builds on our previous observations but also corroborates the prevailing view that successful memory recall depends on the selective reactivation of engram cells^94–97^.

## Discussion

This study demonstrates that SD selectively impairs memory retrievability by reducing the reactivation of DG engram cells without detectable alterations in global DG activity at the time of retrieval. Pharmacological enhancement of cGMP–PKG signaling using vardenafil temporarily restored memory retrieval by supporting the reactivation of DG engram cells. Combining vardenafil treatment with optogenetic engram reactivation resulted in more persistent access of OLMs formed under SD conditions, highlighting the role of engram cells’ reactivation in memory reconsolidation. Whole-brain analysis revealed that SD-induced retrievability deficits extend beyond the hippocampus, impacting downstream cortical and subcortical regions critical for associative memory, which depend on hippocampal output to support effective recall.

### Vardenafil treatment: facilitation of memory recall and reconsolidation

Enhancement of cGMP–PKG signaling using the PDE5 inhibitor vardenafil, administered immediately before testing, effectively restored memory retrieval in SD mice. PDE5 normally degrades cGMP and regulates the Nitric Oxide (NO)–cGMP/PKG pathway, which induces cAMP response element-binding protein (CREB) phosphorylation and supports CREB-dependent LTP^99–101^. By preventing cGMP breakdown, vardenafil elevates intracellular cGMP levels and promotes PKG activation, a signaling cascade known to enhance synaptic efficacy through AMPA receptor trafficking, specifically the phosphorylation and membrane insertion of GluA1- and GluA2-containing subunits^102^. Notably, these mechanistic insights come from studies in which vardenafil was administered during memory consolidation. The role of cGMP–PKG signaling during memory retrieval remains largely unexplored. Nonetheless, it is plausible (but untested) that these cGMP–PKG–dependent processes might contribute to recall. In our study, the vardenafil-mediated memory rescue was accompanied by the selective reactivation of DG engram cells but not to a global increase in DG activity, highlighting a mechanism for reinstating access to stored memory traces. This finding aligns with earlier work showing that memories formed under SD conditions remain latent and recoverable through optogenetic engram reactivation or pharmacological enhancement of intracellular signaling^28^, including modulation of cAMP–PKA signaling via PDE4 inhibition^28^ and protection of cGMP–PKG–dependent processes during SD^42^. However, the effect of acute vardenafil treatment was only temporary, since it did not support successful memory recall days after administration. Therefore, to achieve a more enduring recovery of memory accessibility, vardenafil administration was combined with the optogenetic reactivation of memory engram cells. Vardenafil was administered immediately following optogenetic engram reactivation, viewing the optogenetic manipulation as an artificial re-learning episode. This strategy, which targets the reconsolidation-like processes, resulted in a sustained, manipulation-free restoration of memory recall days later. This result is consistent with our previous study using the combined approach to rescue memories vulnerable to SD^28^. Importantly, the efficacy of this combined treatment seems to relate to the selective re-engagement of memory-specific engram cells, further underscoring that the benefit was not due to a global increase in neuronal activity (as evidenced by the lack of DG total cFos activity changes, Fig.4C-F).

These findings align with the well-established notion that amnesia-like behavioral impairments typically arise from a transient inability to access stored memory traces, rather than from memory erasure. Specifically, such deficits are thought to reflect a failure to reactivate the neuronal ensembles representing the original experience^97^. The observed reduction in DG engram cell reactivation during recall following SD is, therefore, highly consistent with the framework suggesting that successful retrieval depends on the re-engagement of the neuronal population active during initial encoding^103^. Given these considerations, a plausible explanation for the effects of the combinatorial approach during memory reconsolidation is that cGMP-PKG–mediated facilitation of synaptic receptor trafficking temporarily compensates for SD-induced consolidation deficits. This may restore sufficient neuronal excitability to enable selective DG engram reactivation during memory recall several days later. Altogether, these findings position PDE5 inhibition as a useful strategy to facilitate engram-specific mechanisms that restore memory accessibility after SD. Moreover, they strengthen the interpretation that pharmacological enhancement of cyclic nucleotide signaling restores function in compromised, rather than optimal, neural circuits, consistent with ours and other results showing enhancement beyond baseline performance only in SD animals^40,42,104^.

### Distributed cortical–hippocampal network dysfunction after SD

A major novelty of this study is the employment of whole-brain cFos mapping, which provided an unprecedented, system-level view of the network deficits during OLM recall induced by SD. These analyses revealed that successful OLM recall is associated with the integrated engagement of a distributed memory-associative network encompassing both cortical and subcortical regions. This network engagement was disrupted in mice whose memories were rendered inaccessible by SD, as evidenced by significant reductions in cFos density across key nodes involved in the circuitry. This suggests that the causes of SD-induced amnesia are not confined to the hippocampus but instead reflect a disrupted network alteration within the memory-associative circuit^11^. The hippocampus is situated at the core of this network, forming hierarchical circuits with the medial prefrontal cortex, entorhinal and perirhinal cortices, the retrosplenial cortex, and thalamic and subcortical structures^65,105–108^. Interestingly, mice with impaired retrieval of OLMs exhibited reduced activation of the retrosplenial (RSP) cortex. This finding is consistent with a growing body of evidence demonstrating the RSP’s essential role in integrating object and spatial information^83–85^. Moreover, the observation of attenuated activation in the anterior cingulate cortex and entorhinal cortex provides additional evidence that SD broadly perturbs functional integrity across the memory-associative network. The anterior cingulate cortex contributes to storing long-lasting object–place representations, with neural activity patterns persisting over extended delays^86,87^, while the entorhinal cortex, particularly its lateral subdivision, plays a causal role in encoding and recalling object–location associations^109,110^. Collectively, these findings suggest that SD makes OLM memories inaccessible during recall by impairing the coordinated activation of the distributed cortical–hippocampal network required for accurate retrieval of associative representations. Our findings further support the critical role of this network and identify the hippocampal formation as a central hub linking most retrieval-associated structures (T30SS; Fig. 2). Consistent with this, prior studies have shown that disruption of hippocampal output pathways selectively impairs memory retrieval, highlighting the essential role of hippocampal projections in coordinating downstream memory-related circuits^106,108^. Despite not emerging as the most differentially activated regions by cFos, hippocampal output structures (including ProS, SUB, ENTl, and ENTm) exhibited extensive connectivity to nearly all T30SS. This architecture positions the hippocampal formation as a systems-level orchestrator capable of coordinating retrieval-related activity across both cortical and subcortical targets. The apparent dissociation between connectivity and overall cFos expression highlights an important aspect of network organization: regions that exert strong modulatory control over network dynamics may not exhibit large changes in immediate early gene expression. We propose that within these “silent orchestrators” (typically harboring memory engrams), the relevant variable is not total cFos density that matters, but rather the specific engram reactivation. In the context of SD, subtle disruptions in these regions could lead to downstream hypoactivation and retrieval failure without necessitating detectable changes within themselves. Within this framework, the DG emerges as a compelling candidate for an upstream bottleneck. By filtering information entering the hippocampus^111^, it plays a central role in learning, memory, spatial coding, and recall^93,112–115^. Moreover, recent evidence indicates that bidirectional signaling between the DG and ENT is required to preserve synaptic memory engrams^93^. Disruption of this mechanism may therefore underlie impaired engram maintenance and the resulting deficits in memory recall following SD. This mechanism aligns with the concept that successful memory recall depends on the selective reactivation of specific memory engram cells distributed across interconnected brain regions^13,103,116–119^. Consistent with this, using engram-based approaches to selectively reactivate tagged memory traces, we found that object–location information remains stored in the brain even when encoding occurs under sleep-deprived conditions. However, SD impaired the natural reactivation of DG engram cells during recall, resulting in retrieval failure. Notably, pharmacological treatment with vardenafil (either alone or in combination with optogenetic stimulation) restored the reactivation of DG engram cells and rescued memory retrieval. These findings indicate that SD-induced amnesia reflects a disruption in the reactivation of DG memory engram rather than a loss of the stored memory trace itself. Given that engram ensembles are distributed across multiple regions of the altered memory-associative circuit, an important question for future work is whether similar SD-induced disruptions and pharmacological rescue mechanisms occur in other sparse engram ensembles throughout this circuit.

### Translational implications and future directions

Collectively, these findings underscore vardenafil as a promising strategy to mitigate SD–induced memory impairments, with potential relevance to other conditions characterized by deficits in memory retrieval, including Alzheimer’s disease and infantile amnesia. As an FDA-approved PDE5 inhibitor with a well-established safety profile and robust pro-cognitive effects in preclinical models (including both sexes and aged animals) vardenafil provides a strong rationale for clinical translation in memory-related disorders^41^. Nonetheless, several important questions remain. Future investigations should explore whether vardenafil’s restorative effects extend beyond the hippocampus to reestablish functional connectivity among distributed network hubs and rescue engram reactivation in associative regions important for memory recall such as the RSP, ENT, ACA and VIS, thereby advancing our understanding of network-level mechanisms of action. Moreover, the precise molecular mechanisms underlying vardenafil’s pro-cognitive effects remain to be fully elucidated. For instance, it remains unclear whether its facilitation of memory retrieval requires downstream engagement of the cAMP–PKA signaling pathway (as for the consolidation period) or other intracellular cascades beyond the primary cGMP–PKG axis. Furthermore, the long-term stability of memory traces rescued by the combined approach warrant additional study, particularly to determine whether alternative or compensatory circuits are recruited relative to natural recall. Finally, given its encouraging outcomes in preclinical studies and potential relevance to Alzheimer’s disease^99,101,120–122^, future research should rigorously assess the durability, generalizability, and translational potential of vardenafil across diverse memory paradigms and pathological populations. Overall, this study positions vardenafil as a compelling therapeutic candidate to restore memory function following SD and may lay the groundwork for the development of new therapeutic approaches to treat the cognitive deficits that are a debilitating component of many neurocognitive disorders associated with sleep loss and sleep disturbances.

## Acknowledgments

The authors thank Roeland Lokhorst (Microscopy Facility, Netherlands Institute for Neuroscience, Amsterdam, The Netherlands) and Sébastien Courrier (Ingénieur d’étude, Équipe Myologie Translationnelle; Responsable Plateforme Imagerie, INSERM U1251, Aix-Marseille Université, Faculté de Médecine Timone, Marseille, France) for expert assistance with image acquisition. We also acknowledge the veterinary staff and animal care personnel of the University of Groningen animal facility for their support with animal breeding and care. We further thank all current and former members of the Havekes and Silva laboratories for their valuable feedback on the project.

## Lead contact

Further information and requests should be directed to and will be fulfilled by the lead contact, Robbert Havekes

## Author contributions

Conceptualization: CP, PM, RH, BAS

Methodology: CP, CC, AS, RH, PM, BAS, EK, PRAH

Investigation: CP, CC, DMP, ELM, OVH, EYGC, SW, NDV, LMR

Visualization: CP, CC

Funding acquisition: RH, BAS

Project administration: RH, PM

Supervision: PM, RH, BAS

Writing – original draft: CP, CC

Writing – review & editing: CP, CC, DMP, RH, PM, BAS, PRAH, EK

