## Supplementary material for "The phosphodiesterase-5 inhibitor vardenafil reverses sleep deprivation-induced amnesia in mice": S1, S2, S3, S4, S5, S6, S7

### Supplementary information

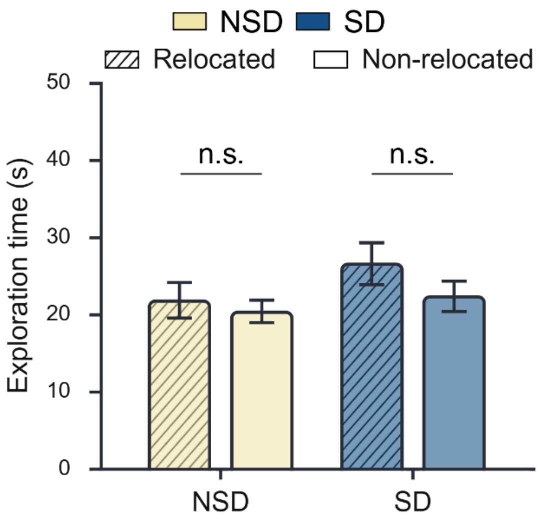


**Figure S1. Exploration times for relocated and non-relocated objects during training corresponding to the experiment shown in Fig. 1.**
No significant differences in object exploration were observed between conditions across any of the three experiments.

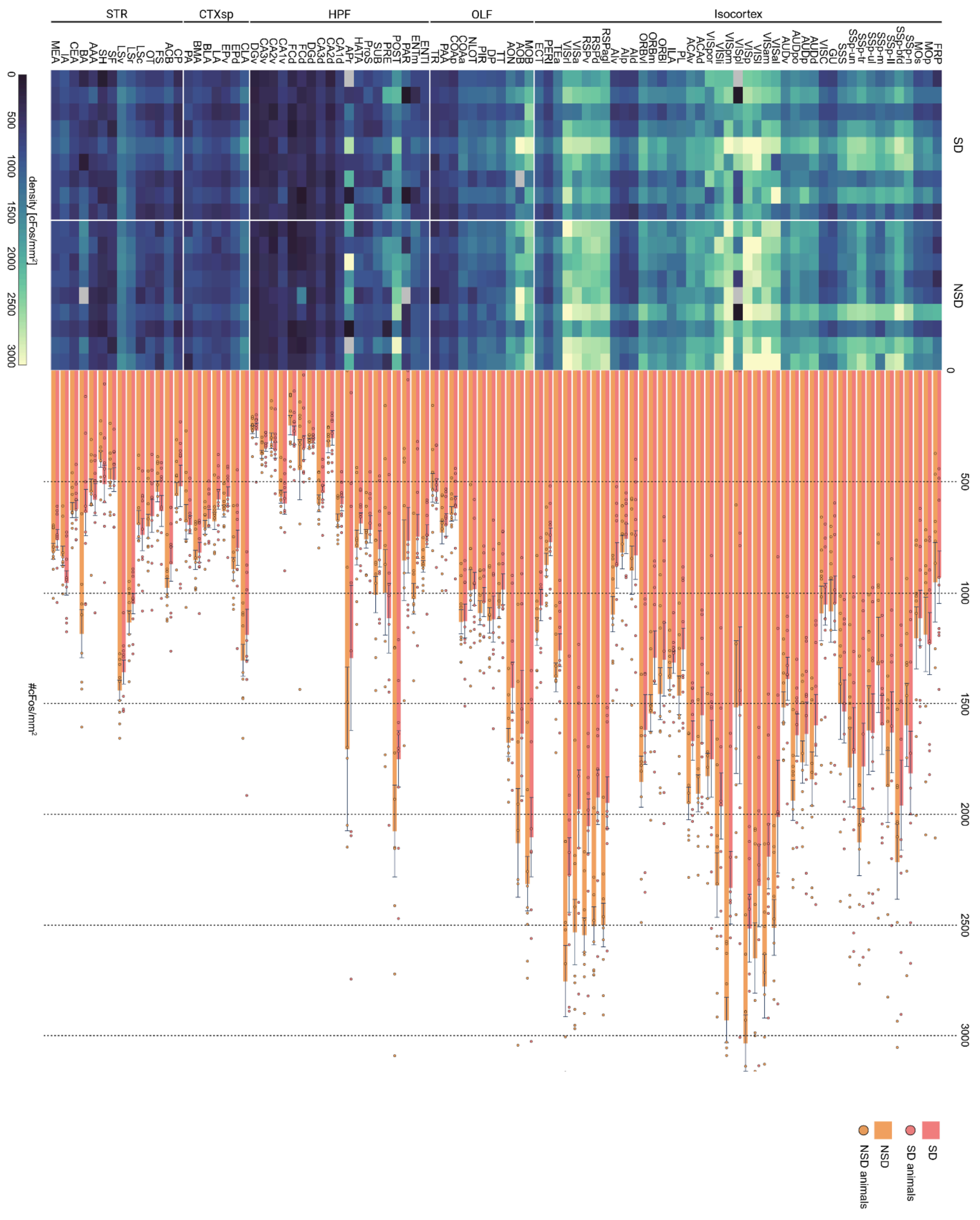


**Figure S2. XmasTree plot of cFos densities in each "summary structure" categorized by CCFv3 major divisions, related to Figure 1.**
Region-by-region cFos densities in Isocortex, olfactory areas (OLF), hippocampal formation (HPF), cortical subplate (CTXsp), striatum (STR), for each animal in each behavioral condition (SD, pink, NSD, orange). Top, heatmaps representing densities for each animal (column) for each region (rows). Gray squares indicate regions that were missing in the image dataset. Bottom, bar plots (mean ± SEM) of cFos densities in NSD and SD groups across all analyzed brain regions (n=9 animals per group).

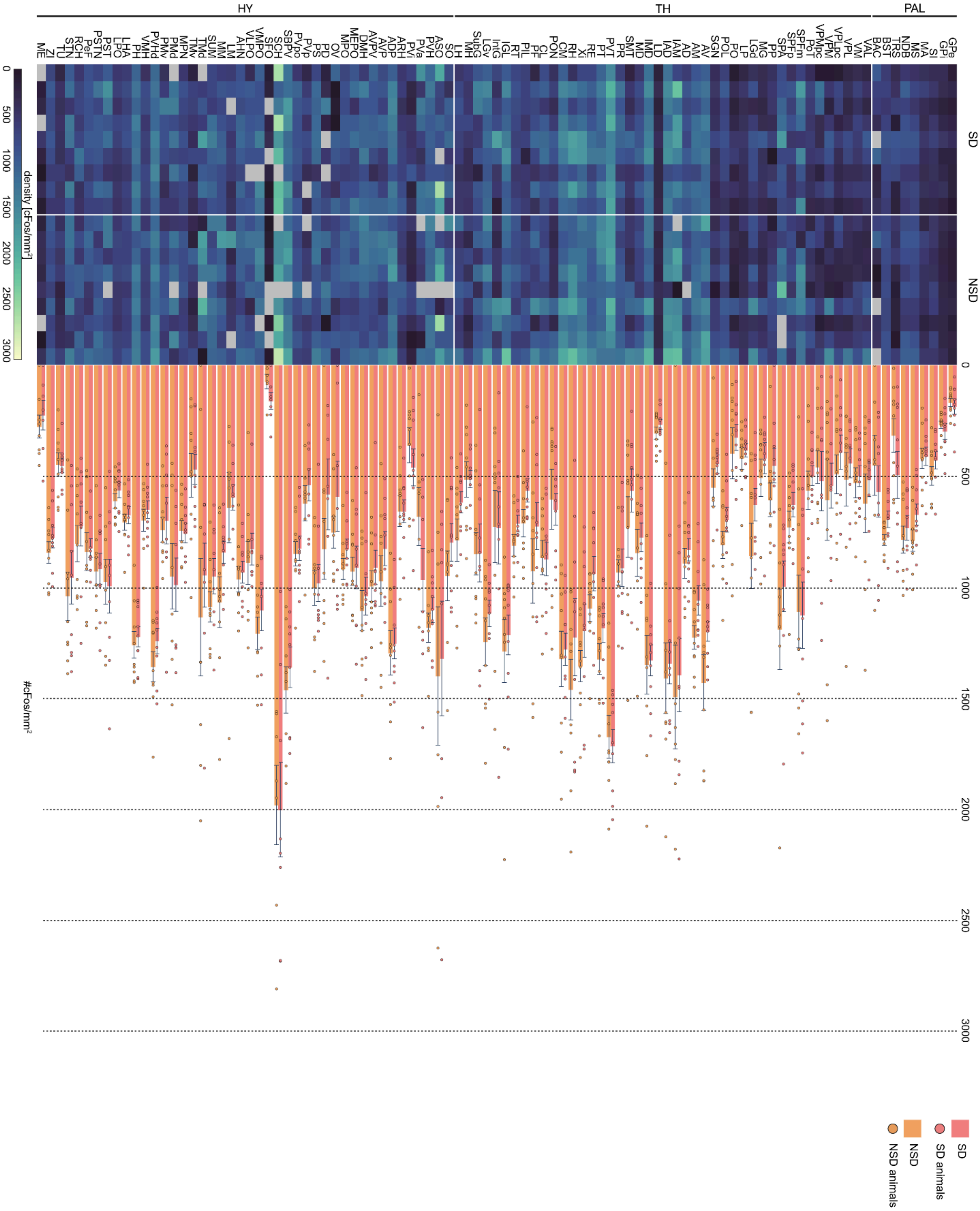


**Figure S3. XmasTree plot of cFos densities in each “summary structure" categorized by CCFv3 major divisions, related to Figure 1.**

Region-by-region cFos densities in Pallidum (PAL), thalamus (TH), hypothalamus (HY), for each animal in each behavioral condition (SD, pink, NSD, orange). Top, heatmaps representing densities for each animal (column) for each region (rows). Gray squares indicate regions that were missing in the image dataset. Bottom, bar plots (mean ± SEM) of cFos densities in NSD and SD groups across all analyzed brain regions (n=9 per condition).

**b**

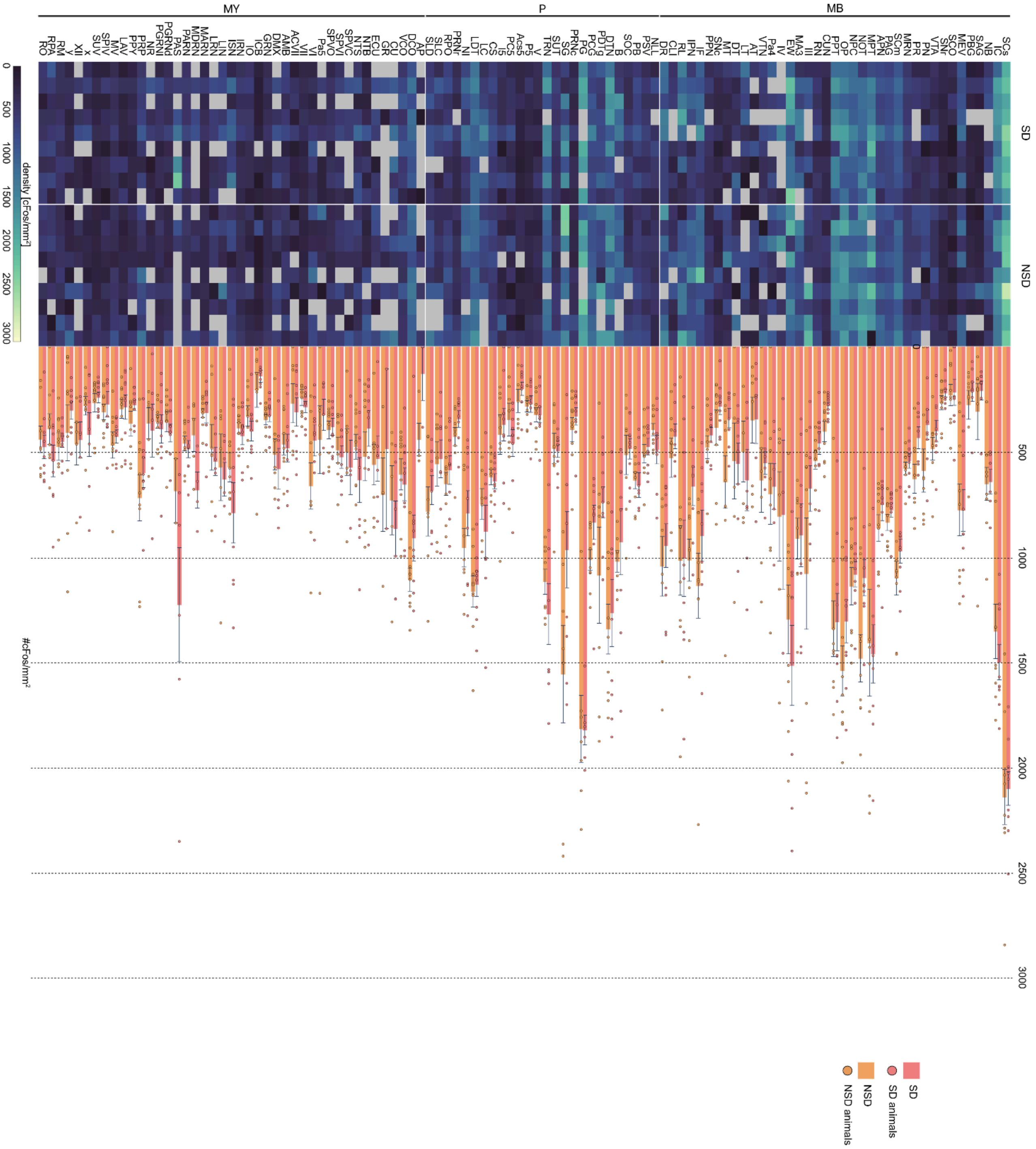

MY
P
MB
SD
NSD
density [cFos/mm²]
0
500
1000
1500
2000
2500
3000
#cFos/mm²
SD
SD animals
NSD
NSD animals

**Figure S4. XmasTree plot of cFos densities in each "summary structure" categorized by CCFv3 major divisions, related to Figure 1.**
Region-by-region cFos densities in Midbrain (MB), Pons (P) and Medulla (MY), for each animal in each behavioral condition (SD, pink, NSD, orange). Top, heatmaps representing densities for each animal (column) for each region (rows). Gray squares indicate regions that were missing in the image dataset. Bottom, bar plots (mean ± SEM) of cFos densities in NSD and SD groups across all analyzed brain regions (n=9 per condition).

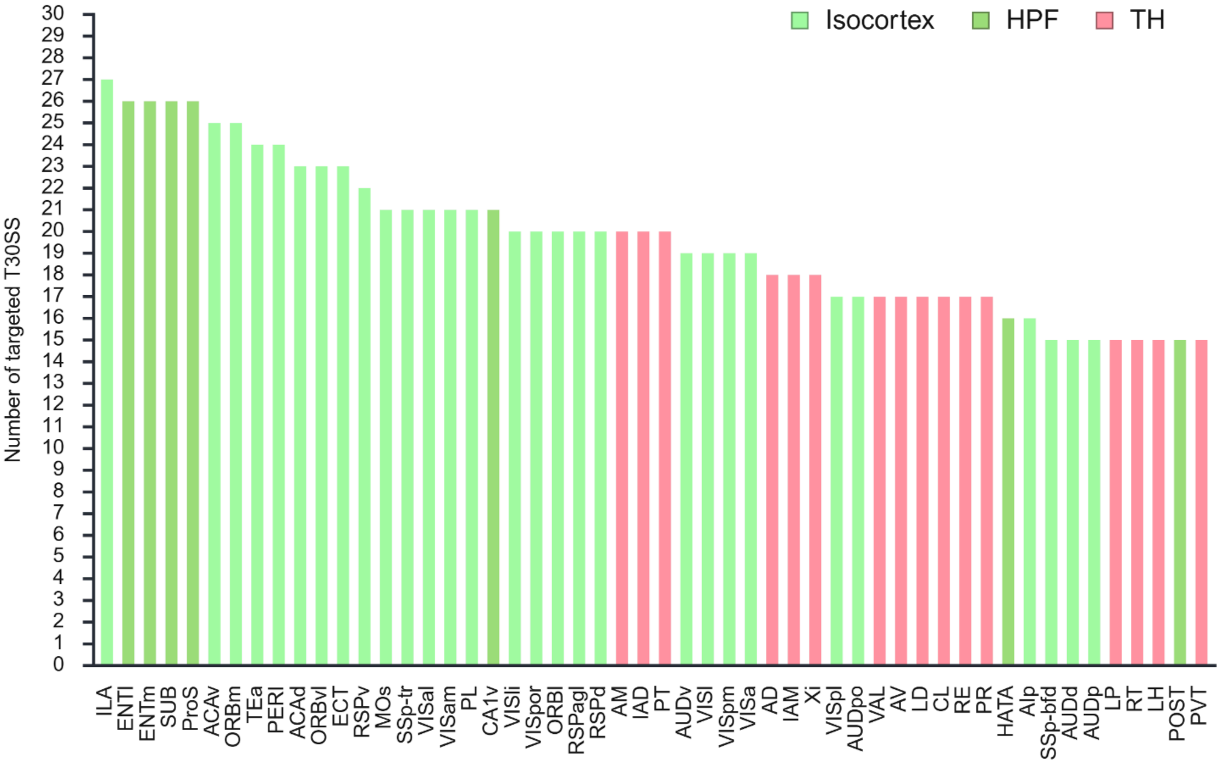


**Figure S5. Rankings of the summary structures projecting T30SS.** Number of T30SS targeted by each summary structure, according to the Allen Institute's data-driven mouse connectome, binarised as explained in Materials & Methods. Each bar is colored based on the major division the corresponding region it belongs to. Regions projecting to less than 15 regions of the T30SS are not displayed; for the complete list, see Table S3.

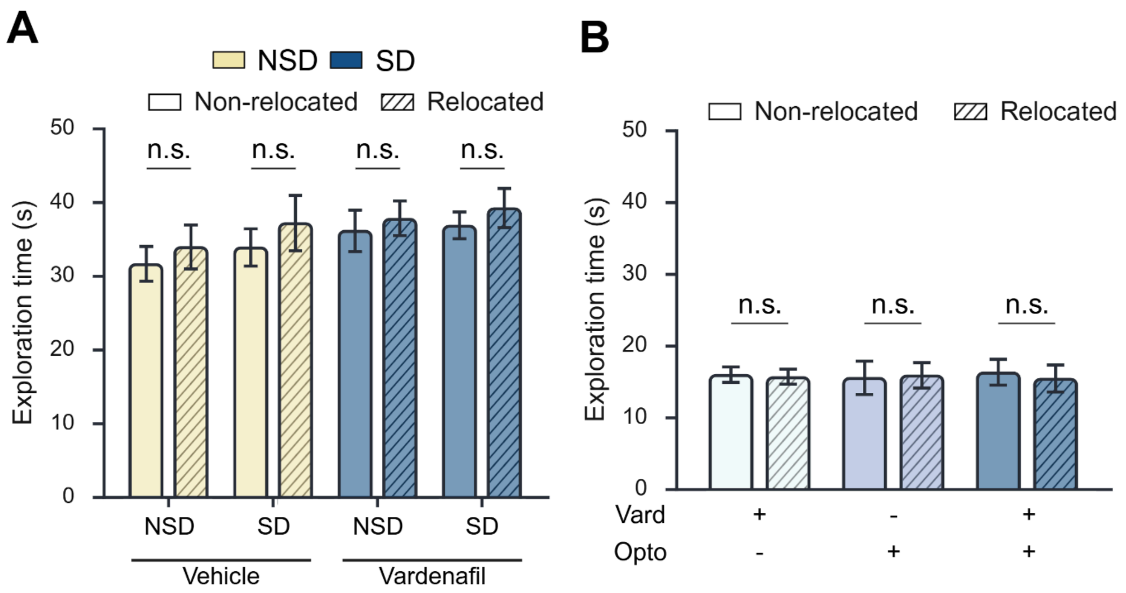


**Figure S6. Exploration times for relocated and non-relocated objects during training.**
Panels (**A–B**) correspond to experiments shown in Figs. 3, and 4, respectively. No significant differences in object exploration were observed between conditions across any of the three experiments.

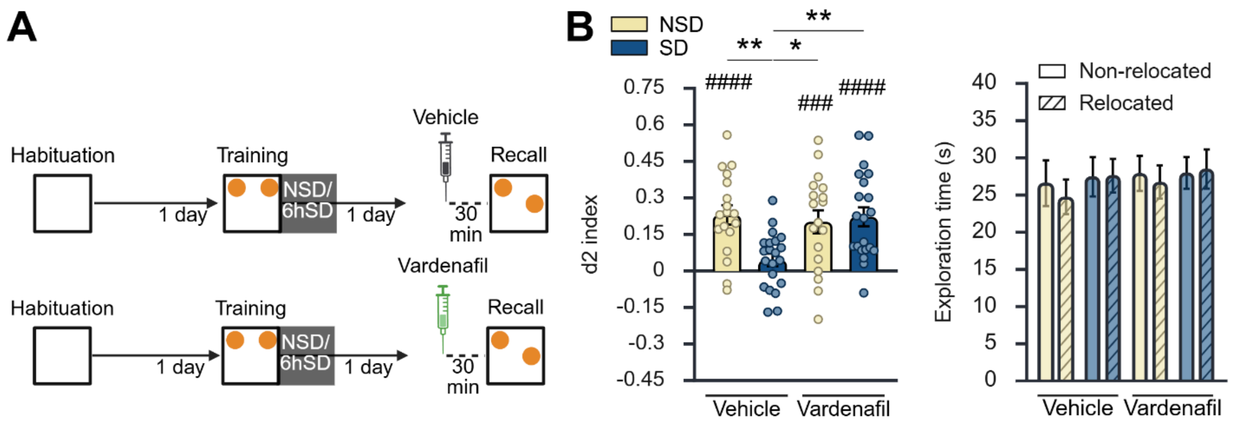


**Figure S7. Vardenafil treatment before testing rescues memory deficits caused by sleep deprivation following training in C57BL/6 male mice.**
**(A)** Wild-type C57BL/6 male mice (n = 18 per condition) were first habituated to the empty arena. After 24 hours, they were subjected to 6 hours of sleep deprivation (SD) or left undisturbed (NSD) immediately after training in the OLM task. Animals receive Vardenafil (0.3 mg/kg, i.p. injection) or vehicle 30 min before the retention test (24 hours post-training). **(B)** Left: d2 discrimination index. As expected, NSD groups scored significantly above chance level, indicating proper detection of spatial novelty and successful memory retrieval, whereas vehicle-treated SD mice failed to detect spatial novelty. Importantly, SD mice treated with vardenafil 30 minutes before testing demonstrated proper detection of spatial novelty, comparable to NSD groups [significant interaction: p = 0.008; post-hoc: p=0.002 for NSD-vehicle vs. SD-vehicle, p=0.026 for NSD-vardenafil vs. SD-vehicle, p=0.0005 for SD-vardenafil vs. SD-vehicle]. Right: Exploration time of the two objects during the training trial. The 'non-relocated' object stays in the same location

during the subsequent test trial, while the 'relocated' object is moved to a new location. Mice across all experimental conditions showed no significant object preference during training, spending similar amounts of time exploring both objects [N.S.]. The dependent variable was $log_{10}$-transformed before analysis to meet the assumption of homogeneity of variances. Data are presented as mean ± SEM. Statistical significance is denoted as */#p<0.05, **/##p<0.01, ***/###p<0.001, ****/####p<0.0001.  # indicates significantly different from zero.
