## supplementary materials for "The phosphodiesterase-5 inhibitor vardenafil reverses sleep deprivation-induced amnesia in mice"

### An extended description of all behavioral experiments

*Whole-brain differences between accessible and inaccessible memories (Figure1)*
To investigate whole-brain differences between accessible and inaccessible memories, we exposed adult male C57Bl/6J mice to an OLM training task. Mice were then either sleep deprived for six hours (SD group) or left undisturbed (NSD group). The day after, mice were tested in the OLM task, as previously described. Ninety minutes after the test trial, mice were euthanized by transcardial perfusion.

*Treatment with vardenafil 24 hours after training, 30 minutes prior to the retrieval test (Figure 3 and S7).*
The vardenafil treatment, 30 minutes prior to the test, 24 hours after training was performed on c-fos-tTA mice bilaterally injected with AVV9-TRE-ChR2-mCherry virus and put on a Dox chow (40 ppm). After at least three weeks from viral surgery, mice were handled and habituated as previously described. Following habituation, they were switched to regular chow to enable engram labelling to occur during the next day's training. Immediately after training, mice were put on 1000 ppm Dox chow. Half of the group were SD for 6 hours (SD group), while the other half were left undisturbed (non-sleep deprived, NSD group). Twenty-four hours after training, mice received an i.p. injection of vardenafil (dose: 0.3 mg/kg) or vehicle solution 30 minutes before testing. Ninety minutes after the test trial, mice were sacrificed by means of transcardial perfusion. The same was done in adult male C57Bl/6J mice (Fig.S7), however here we followed a two-by-two factorial design with 'group' (sleep deprived/non-sleep deprived) as a between-subject factor, 'treatment' (vardenafil/control) as a within-subject factor, therefore mice were reused once to ensure statistical power while complying with the animal welfare principles of reduction.

*Optogenetic engram reactivation in combination with vardenafil treatment (Figure 4)*
Similar to previously described studies, mice were first handled and habituated. At the end of the third habituation day and 24 hours before the training trial, mice were taken off the Dox food. On experimental day 1, mice underwent the learning trial in which they were allowed to explore two similar objects for 10 min. After the training trial, we immediately switched the animals to a 1000 ppm Dox diet and subjected all of them to 6 hours of SD. The animals were then left undisturbed for the following three days. On day 4, the reactivation session took place, and mice were divided into three experimental conditions: 'laser on + vehicle', 'laser off + vardenafil, and 'laser on + vardenafil'. The reactivation took place in the home cage and lasted 3 minutes (as described above). During this 3-minute period, the laser was turned on in the 'laser on' experimental condition, while in the "laser off" group, mice were only attached to the laser cables but received no stimulation. Immediately after reactivation, we i.p. injected the mice with either vardenafil or vehicle solution. Two days later, mice were subjected to the testing session in which one of the two objects was moved to a novel location. During testing, no drugs or laser treatment were applied. Ninety minutes after the test trial, mice were euthanized by transcardial perfusion.
